# Characterization of the *Grin1*^Q536R/+^ mouse: a preclinical model for *GRIN1*-related neurodevelopmental disorder

**DOI:** 10.64898/2026.09.09.750465

**Authors:** Megan T. Sullivan, Yuanye Yan, Lyra Vania, Patrick Tidball, Quinn Pauli, Tatiana V. Lipina, Sridevi Venkatesan, Peter S.B. Finnie, Emily Fujiwara, Wendy Horsfall, John Georgiou, Robert E. McCullumsmith, Evelyn K. Lambe, Robert P. Bonin, Ali Salahpour, Graham L. Collingridge, Amy J. Ramsey

## Abstract

N-methyl-D-aspartate receptors (NMDARs) are ionotropic glutamate receptors playing critical roles in brain development, synaptic plasticity, and cognition. *GRIN1*-Related Neurodevelopmental Disorder (*GRIN1*-NDD) is a rare genetic condition caused by pathogenic variations in the *GRIN1* gene, which encodes the obligatory subunit of NMDARs. The spectrum of *GRIN1* clinical symptoms is hypothesized to result from the functional consequences that different missense variants have on NMDARs. To investigate the disease mechanism *in vivo*, we generated a novel heterozygous *Grin1*^Q536R/+^ knock-in mouse model that carries the identical variation as an adolescent male patient. We describe the clinical presentation of this patient and conduct comprehensive molecular, morphological, electrophysiological, and behavioural characterization in the juvenile, adult, and aging mice. Compared to wildtype littermates, *Grin1*^Q536R/+^ mice displayed reduced whole-cell NMDA-induced currents in cortical pyramidal neurons, and reduced NMDAR-mediated synaptic transmission, decreased long-term potentiation, but intact depotentiation at the hippocampal CA1 synapses. Morphological change was observed in the dentate gyrus region of the *Grin1*^Q536R/+^ mice. Behavioral phenotyping revealed age- and sex-dependent differences from controls, including hyperlocomotion, reduced muscle strength, and spatial learning deficits. These phenotypes are in line with the clinical manifestations and the relative disease severity of the male patient. The age-dependent phenotypic shift in *Grin1*^Q536R/+^ mice highlights the model’s value for investigating *GRIN1*-NDD disease progression and informing longitudinal monitoring as well as potential therapeutic adjustments with age. Taken together, our findings establish a novel and robust in vivo model for studying NMDAR mechanisms and disease pathology in *GRIN1*-NDD, while supporting the preclinical development of new therapeutic strategies.

**Significance statement:** Based on an adolescent male patient carrying a *de novo* heterozygous Q536R variant in the *GRIN1* gene, we report the creation and comprehensive molecular, electrophysiological and behavioural characterization of the *Grin1*^Q536R/+^ mouse model. Our findings reveal that the Q536R/+ variant confers an *in vivo* phenotype indicative of loss of NMDAR function, particularly deficits in NMDAR-mediated transmission and long-term potentiation. *Grin1*^Q536R/+^ mice display neuroanatomical abnormalities and relatively mild behavioural phenotypes, mirroring the observed clinical severity of the patient. The present study contributes a distinct clinically relevant model of *GRIN1* variation that represents aspects of the *GRIN1*-NDD phenotypic spectrum that are poorly studied. Our findings provide insight into disease mechanisms and support the preclinical development of potential therapeutics for *GRIN1*-NDD.

## Introduction

N-methyl-D-aspartate receptors (NMDARs) are tetrameric ionotropic glutamate receptors that play critical roles in synaptic plasticity, brain development, and cognition. Rare, often *de novo* variants in NMDAR-encoding *GRIN* genes have been identified in patients with neurodevelopmental disorders (Li et al., 2019; Benke et al., 2021; Korinek et al., 2024). *GRIN1*- Related Neurodevelopmental Disorder (*GRIN1*-NDD) is characterized by clinically significant variations in the *GRIN1* gene (Platzer and Lemke, 2019), which encodes the obligatory GluN1 subunit of NMDAR (Kleckner and Dingledine, 1988; Clements and Westbrook, 1991; Moriyoshi et al., 1991). *GRIN1*-NDD has an estimated incidence of 5 per 100,000 births (Lemke, 2020). Affected individuals display mild-to-profound developmental delay and intellectual disability. With variable frequency and severity, patients also exhibit symptoms such as epilepsy, muscular hypotonia, movement disorders, etc. (Ohba et al., 2015; Lemke et al., 2016; Platzer and Lemke, 2019).

More than 100 unique pathogenic or likely pathogenic *GRIN1* variants have been identified, most of which are missense variants (Korinek et al., 2024) (https://www.rarevariants.org/grindb/database/). These variants occur throughout the entire GluN1 protein (Lemke et al., 2016; Hansen et al., 2021; Korinek et al., 2024) but are enriched in the agonist binding (ABD) or transmembrane domains (XiangWei et al., 2018; Ragnarsson et al., 2023; Xu et al., 2024), which regulate receptor gating and ion permeation (Amin et al., 2021). One such variant, p.Gln536Arg (Q536R), is in the S1 region of the ABD harbouring the glycine binding site. This variant has been identified in three individuals; the two with available clinical data presented with seizures, mild developmental delay, intellectual disability, and autism (https://grin-portal.lalresearchgroup.org/).

*GRIN1* variants have diverse impacts on NMDAR function, including but not limited to agonist/co-agonist potency, allosteric modulation, channel open probability, response kinetics, and receptor trafficking (Korinek et al., 2024). To establish structure-function relationships and to guide patient treatment, *in vitro* assays have been established to functionally classify *GRIN1* variants as gain- or loss-of-function (GoF or LoF) (Myers et al., 2023). NMDARs containing GluN1-Q536R subunit displayed reduced glutamate potency (3.7-fold), markedly reduced glycine potency (3432-fold), and altered sensitivity to Zn^2+^ and proton inhibition, supporting a LoF classification (CFERV, Emory University). Because extracellular glycine (or d-serine) concentrations are relatively constant, this profound reduction in glycine potency is predicted to limit glycine occupancy and reduce receptor availability for activation (Kew et al., 2000; Hansen et al., 2018).

Mouse models of *GRIN* variation were initially established to investigate receptor function (Mohn et al., 1999; Kew et al., 2000; Single et al., 2000). More recently, patient-specific knock-in mouse models are developed as *in vivo* platforms to elucidate disease pathophysiology and evaluate therapeutic strategies (Benke et al., 2021). Previous *in vitro* studies showed that the position of a variant within *GRIN1* gene, together with its impact on NMDAR function, may serve as predictor of patient outcomes (Cha et al., 2025; Montanucci et al., 2025). Although multiple patient-derived *Grin1* mouse models have been generated (Lee, 2023; Sullivan et al., 2024), no patient-specific mouse model exists for variants within the *Grin1* S1 region. A related model targeting this domain, *Grin1*^D481N^ mouse, was generated to examine the consequences of reduced glycine affinity (Kew et al., 2000). However, this variant does not correspond to any known patient variant and limited studies have since been conducted (Labrie et al., 2008).

We generated a *Grin1*^Q536R/+^ knock-in mouse model carrying the heterozygous Q536R variant identified in a *GRIN1*-NDD patient, the first clinically relevant mouse model for the *GRIN1* S1 region. Although the biophysical properties of this variant have been characterized *in vitro*, its impact on brain physiology and behavior remained unknown. We show that *Grin1*^Q536R/+^ mice exhibit reduced synaptic NMDAR transmission, impaired long-term potentiation, and mild but detectable behavioral phenotypes that parallel the relative mild clinical presentation reported for this variant. By capturing variant-specific mechanisms and phenotypes not presented in previous models, this work introduces a distinct and translationally relevant model for understanding NMDAR function, dissecting *GRIN1*-NDD pathophysiology, and supporting preclinical therapeutic development.

## Materials and Methods

### Collection of Patient Information

Consent was obtained to share de-identified patient data with collaborating researchers under a protocol approved by the University of Toronto Research Ethics Board (#00047662; Use of patient medical information in the study of *GRIN1*-Related Neurodevelopmental Disorder).

Upon provision of written informed consent, the caregiver of the individual carrying the Q536R variant was provided with a questionnaire. The questionnaire consisted of five sections: General information, Diagnosis, Current Symptomology & Treatment Regimens, Previously Tried Medications and Personal information. Questions inquired about patient age, sex, clinical presentation at birth, the process of receiving a *GRIN1*-Related Neurodevelopmental Disorder diagnosis, symptomology, current treatment regimens and responses to previously tried medications, as well as the patient’s strengths, challenges and unmet treatment needs.

### Animals

*Grin1*^Q536R/+^ mice were generated by CRISPR-Cas9 endonuclease-mediated transgenesis via pronuclear injection and identified by Sanger sequencing. Heterozygous *Grin1*^Q536R/+^ mice were crossed with C57BL/6J mice to produce *Grin1^+/+^* (wildtype or WT) and *Grin1*^Q536R/+^ (Q536R/+) mice which were used for experimentation. Genotypes of experimental mice were confirmed by two separate PCR reactions for each allele: the WT and Q536R variant allele (different by 2 base pairs). A control PCR (thrombomodulin) was multiplexed in each PCR reaction to ensure true negative result (see supplementary materials for primers). Touch down PCR was performed as previously described (Sullivan et al., 2024). PCR products were checked and visualized by gel electrophoresis.

All experimental mice were group housed on a 12-hour light-dark cycle with *ad libitum* access to food and water. All procedures were conducted during the light phase and complied with animal use protocols and guidelines of the University of Toronto Temerty Faculty of Medicine and Pharmacy Animal Care Committee, The Center for Phenogenomics’ Animal Care Committee, and Canadian Council on Animal Care.

### RNA isolation and RT-qPCR

Mice were anesthetized with isoflurane and euthanized by cervical dislocation. Forebrain tissue was dissected and homogenized in TRIzol reagent (Invitrogen). RNA was isolated by chloroform phase, precipitated in isopropanol, washed by 70% ethanol and resuspended in ultrapure water. Isolated RNA samples were immediately reverse transcribed to synthesize cDNA using SuperScript VILO IV Master Mix with ezDNase Enzyme (Invitrogen) following manufacturer’s manual. qPCR was performed in 20μL volume using Power SYBR Green Master Mix with QuantStudio 3 Real-Time PCR system (Applied Biosystems).

### Synaptic plasma membrane isolation and western blots

Mice were anesthetized with isoflurane and euthanized by cervical dislocation. The anterior third of the brain tissue was coronally dissected and homogenized in 0.32M HEPES-buffered Sucrose (containing protease/phosphatase inhibitor cocktail) to obtain total protein lysate samples. Synaptic plasma membrane fraction was isolated from the total protein lysates as previously described (Bermejo et al., 2014; Sullivan et al., 2024). Loading samples were prepared in NuPage LDS sample buffer (Invitrogen), 5% β-mercaptoethanol and heated to 55℃ for 15min. Protein samples (10μg) were electrophoresed in a 4-12% Bis-Tris gel (Thermo Fisher Scientific) using MOPS running buffer (50mM MOPS, 50mM Tris, 1mM EDTA, 0.1% SDS), which was then transferred onto immobilon PVDF membrane (Millipore). Blots were blocked in 3% skim milk dissolved in TBST (Tris-buffered saline with 0.1% Tween 20). Primary antibodies used: rabbit anti-NMDAR1 (1:1000, AB9864R, Sigma-Aldrich), rabbit anti-NMDAR2A (1:1000, AB1555P, Sigma-Aldrich), rabbit anti-NMDAR2B (1:1000, AB1557P, Sigma-Aldrich). Secondary antibody used: IRDye 800CW goat anti-rabbit (1:5000, 925-32211, LI-COR Biosciences). Blots were imaged using Odyssey M Imager and normalized to REVERT 700 total protein stain (LI-COR Biosciences).

### Cortical whole-cell electrophysiology

Whole-cell patch-clamp recordings were performed on layer 5 pyramidal neurons in 400 µm thick medial prefrontal cortex (mPFC) brain slices from adult *Grin1*^Q536R/+^ and wildtype (+/+) littermate mice as previously described (Venkatesan et al., 2023, 2026) and summarized here: whole-cell currents evoked by bath application of NMDA (20 μM, 30s) were recorded under voltage-clamp at -75 mV, using a K-gluconate patch solution containing (in mM) 120 potassium gluconate, 5 KCl, 10 HEPES, 2 MgCl_2_, 4 K_2_-ATP, 0.4 Na_2_-GTP, and 10 sodium phosphocreatine, adjusted to pH 7.3. A modified artificial cerebrospinal fluid (ACSF) containing low Mg^2+^ (0.5 mM MgSO_4_) and 5mM KCl was used to reduce voltage-dependent Mg^2+^ blockade of NMDARs. AMPA receptor-mediated currents were blocked with 20 μM CNQX.

### Hippocampal slice field potential electrophysiology

Dorsal hippocampal slices prepared from adult (postnatal week 10–16) male and female Q536R/+ and wildtype (+/+) mice were used for electrophysiological recordings. Experimenters were blinded to genotype, and separate experiments were performed to assess 1) basal AMPAR- and NMDAR-mediated synaptic transmission, and long-term potentiation (LTP), and 2) depotentiation of LTP. The first study was performed as previously described (Sullivan et al., 2024) and depotentiation of LTP was performed as described below.

#### Depotentiation of long-term potentiation

Mice were first anesthetized with 2.5% Avertin (Tribromoethanol) and euthanized by trans- cardiac perfusion with a cold (2-4^0^C) sucrose-based solution containing (in mM): 50 Sucrose, 92 NaCl, 15 Glucose, 5 KCl, 1.4 NaH_2_PO_4_, 26 NaHCO_3_, 0.5 CaCl_2_, 7 MgCl_2_ and 1 Kynurenic acid (osmolality 322-330 mOsm, bubbled with 95% O_2_ and 5% CO_2_). 400 µm sagittal slices were cut in ice cold ACSF containing (in mM): 124 NaCl, 10 glucose, 26 NaHCO_3_, 3 KCl, 1.4 NaH_2_PO4, 1 MgSO_4_ and 2 CaCl_2_ (osmolality 300-310 mOsm, bubbled with 95% O_2_ and 5% CO_2_) using a Leica VT1200 S vibratome (Leica Biosystems, Deer Park, Illinois, United States). The CA3 region was carefully dissected from each slice before recovery in 28^0^C ACSF bubbled with 95% O_2_ and 5% CO_2_ for at least 1.5 hours prior to recording.

Field excitatory postsynaptic potentials (fEPSPs) were recorded at Schaffer collateral-CA1 synapses in a submerged-type recording chamber perfused with oxygenated ACSF maintained at 28.5^0^C (± 0.5^0^C) at a rate of 1.5 mL/min. fEPSPs were evoked every 30 seconds (0.033 Hz) using a bipolar platinum-iridium stimulating electrode (cat nr. #30250, FHC, Bowdoin, Maine, United States) controlled by a constant current stimulus isolator (A365, World Precision Instruments, Sarasota, Florida, United States) (0.1 ms pulse width) and recorded with an ACSF- filled borosilicate glass micropipette (1-3 MΩ). Signals were amplified using a Multiclamp 700A amplifier (Axon Instruments, Scottsdale, Arizona, United States) and low pass filtered at 2.4 kHz and digitized at a sampling rate of 20 kHz with an Axon Digidata 1550B (Axon Instruments). Stimulus strength was adjusted to maintain half-maximal fEPSP amplitude according to an input/output (I/O) curve. Following stable baseline recordings lasting at least 30 minutes, LTP was induced using either a spaced or compressed theta burst stimulation (TBS) consisting of five bursts at 5Hz, with each burst composed of five pulses at 100Hz and each train repeated 3 times spaced apart by either 10 minutes (spaced TBS) or 10 seconds (compressed TBS) (as in (Park et al., 2016)). To induce depotentiation (DEP), a 2Hz (10 minutes) low-frequency stimulation (LFS) was administered 30 minutes after the last TBS as in (Park et al., 2019). fEPSPs were monitored for 30 minutes after DEP induction. The fEPSP descending slope (mV/ms) was measured from the termination of the fiber volley to the initial peak and expressed as a percentage of the mean baseline response. The percent DEP was calculated by subtracting the average fEPSP 20-25 minutes (sLTP-DEP), or 25-30 minutes (cLTP-DEP) after administration of the LFS from the average fEPSP obtained 10-15 minutes after the final TBS administration. To probe for any accompanying presynaptic alterations following LTP or DEP induction, paired-pulse ratios (PPRs) of fEPSPs were calculated following administration of two pulses delivered 50 ms apart. PPRs were obtained at baseline, 30 minutes after the last TBS administration (LTP) or 30 minutes after LFS administration (post-LFS). Representative traces are an average of 5 traces obtained at baseline, after LTP induction or after DEP induction. All recordings were monitored and analyzed using pClamp v10.7 (Axon Instruments). Electrophysiological recordings were interleaved.

### Histology

Mice were anesthetized with isoflurane and euthanized by cervical dislocation. Whole brains were prepared as stated previously (Sullivan et al., 2024). 5 μm thick sagittal sections that were 1.525 mm lateral to Bregma were stained with hematoxylin and eosin or Nissl stain. Stained brain sections were imaged using the Axioscan 7 Slide Scanner (Zeiss) and images were analyzed with QuPath v0.3.2 (Bankhead et al., 2017). Thickness and cell layer measurements of the neocortical and hippocampal regions were performed as previously reported (Amador et al., 2020; Edwards et al., 2020; Sullivan et al., 2024). All analyses were conducted under conditions blinded to genotype.

### Behavioural phenotyping

A battery of behavioural tests was performed to measure different domains of behaviour at different ages that are relevant to *GRIN1*-NDD. Righting reflex, ultrasonic vocalization and wire hang tests were performed at early developmental stages (postnatal day 6 and 21, respectively). In adult mice (3-5 months) we conducted wire hang, open field, Y-maze, social behavior, and Barnes maze tests. Older adult mice (9-13 months) were tested with open field, Y-maze, and social behaviour tests. The presence of behavioural convulsions was systematically monitored for 12 weeks starting with mice at 5-7 months of age. Both male and female mice were included in each behavioural test. Separate cohorts of mice were tested to minimize confounds of repeated testing. The tests were performed as summarized below.

#### Maternal isolation induced ultrasonic vocalization (USV) and righting reflex

Maternal isolation induced USV and righting reflex were collected at postnatal day (PND) 6 and PND 21 in mice of both sexes to assess potential developmental delay and/or communication deficits. All data were collected and analyzed as previously described (Coffey et al., 2019; Sullivan et al., 2024).

#### Wire hang test

Wire hang test was performed to assess motor function and muscle strength in both male and female mice at PND 21 and in adulthood (3-5months) as previously described (Sullivan et al., 2024). The holding impulse was calculated as average latency to fall (s) / body weight (kg).

#### Open field test

A 2-hour open field test was performed to assess general locomotor activity and stereotypy of the experimental mice. The test was conducted in a 43.5cm^2^ box with three 16 beam infrared arrays (X, Y, and Z axes) in an isolated sound-attenuating chamber with dim light (30 lux). The experimental arena was created by inserting a divider in the box that creates four equal arenas (∼21 cm^2^). Two sex-matched mice were tested in diagonally adjacent arenas within the same chamber. The mouse was placed in the center of the arena, and the animal activity was tracked for 2 hours with 5min bins, which was analyzed by Activity Monitor Software (Med Associates, Inc.).

#### Y-maze

The Y-maze was performed to assess the spatial working memory as previously described (Mandillo et al., 2008; Milenkovic et al., 2014; Chen et al., 2018). Each mouse was initially placed at the center of the apparatus and allowed to explore all three arms for 5 minutes. Number of entries to each arm and spontaneous alterations were recorded manually with the assessor blinded to the genotypes. Spontaneous alteration behavior was defined as consecutive visits of all three arms. % alteration = number of spontaneous alterations / (total arm entries – 2) *100%.

#### Sociability and social novelty preference

Social interaction behaviours were tested in a rectangular, one-chamber box (59 x 39 x 22 cm). Two identical cylindrical mouse enclosures (diameter: 11 cm, height: 15 cm) with grid bars (7 mm apart) were placed in the opposite corners of the rectangular chamber. Sex- and age-matched naïve wildtype mice that were socially novel to the experimental mice were used as social-partner mouse in this test. The experimental mouse was placed in the center of the chamber (containing two empty enclosures) and left to explore for 5 minutes (habituation). Then, a social-partner mouse was randomly placed in one of the two enclosures. The experimental mouse was then allowed to explore the chamber for 10 minutes (sociability). After the sociability session, the social-partner mouse remained in the same enclosure (familiar mouse), and a second social-partner mouse was placed into the empty enclosure in the opposite corner (novel mouse). The experimental mouse was then allowed to explore the chamber for another 10 minutes (social novelty). The test was video recorded by an overhead camera, which was analyzed by EthoVision XT (Noldus Information Technology, Netherlands). A 5 cm zone around each cylindrical enclosure was defined as the approach zone. The duration of stay and number of visits to the approach zone were automatically measured by EthoVision XT. Time per visit = duration of stay in the approach zone/number of visits to the approach zone.

#### Barnes maze

The Barnes maze was performed to assess spatial learning and memory (Pitts, 2018). The Barnes maze apparatus is a circular platform (diameter: 121 cm) containing 40 equally spaced holes located 2.5 cm from the edge. Spatial cues with distinct patterns and shapes were placed on the walls in the test room as visual cues for spatial orientation. A black goal box (11 x 6 x 6 cm) enriched with home cage bedding and nesting materials was placed under one hole (escape hole or goal zone). The platform was evenly divided into four quadrants with the escape hole located in the center of the holes in one quadrant (the goal quadrant). Aversive stimuli (bright light and loud music) were turned on during the trials. The experiment consisted of three sessions: habituation (day 1), training (day 2-5), and testing (day 5, 5 hours after training). During habituation, mice were placed between the platform center and goal box and left to roam for 3 minutes across 2 trials. Mice were guided into the goal box if failed to enter independently. For all trials in which mice entered the goal box, mice remained in the goal box for 30s before being returned to their home cage. During training sessions, the experimental mouse was placed in the maze and allowed to explore and escape to the goal box with a 3-minute cut off. Each mouse was trained 4 trials per day with different starting locations for each trial (from the center, the right side, the left side, and the opposite side of the escape hole). The training was repeated for four days. On the test day, the goal box was removed, and the mouse was allowed to freely explore the maze for 90 seconds starting from the center of the maze (probe trial, 1 trial per mouse). The experiment was video recorded with an overhead camera. Travel distance, time to escape (primary latency), and time in goal quadrant were automatically analyzed by EthoVision XT (Noldus Information Technology, Netherlands).

### Behavioural Convulsion Tracking

*Grin1*^Q536R/+^ and corresponding wildtype mice (age 22-29 weeks) were monitored once weekly for 12 weeks for the presence of behavioural convulsions and evaluated as previously described (Sullivan et al., 2024). Behavioural convulsion severity was rated by genotype-blinded raters from videos using Racine scoring as depicted in Table S1.

### Experimental Design and Statistical Analyses

Values are presented as mean ± SEM with error bars representing SEM unless otherwise stated. Statistical analysis was conducted using GraphPad Prism version 10.1.1 (GraphPad Software, USA). Unpaired t-test and repeated measures two-way or three-way ANOVA analysis (followed by Tukey’s or Šidák post hoc analysis when appropriate) were performed as indicated in figure legends. For *ex vivo*, molecular, and Barnes maze analyses, sexes were combined for statistical analyses due to limited sample size. Sex differences were analyzed for all other *in vivo* assays, including body weight and behavioural tests. When no significant main effect of sex was detected, data from both male and female mice were combined for graphical presentation. For field potential electrophysiology recordings, I/O curves were derived using simple linear regression analysis of fibre volley (FV) amplitude against stimulus intensity and NMDAR- or AMPAR-mediated fEPSP slopes against FV amplitude, to determine the stimulus-to-FV and FV- to-fEPSP relationships, respectively. Statistical significance was set at p < 0.05.

## Results

### Patient Clinical Characteristics

The patient is a 17-year-old male who initially presented with developmental delay and low muscle tone and was diagnosed with non-syndromic autism spectrum disorder at 8 months. The patient later experienced a visible seizure at 5 years old and has had only one visible seizure since (at 8 years old). Current symptomology includes cortical visual impairment, challenges maintaining attention, poor fine motor skills and behaviours associated with mild intellectual disability and autism spectrum disorder. This includes challenges with age-appropriate social communication and speech. The patient’s level of cognitive functioning permits completion of tasks associated with daily living and school attendance. Thus, this individual could be considered relatively high functioning compared to others on the *GRIN1*-NDD phenotypic spectrum.

The patient’s current pharmaceutical treatment regimen includes atomoxetine for attention difficulties and valproate for seizure control **(Table S2)**.

### Patient Clinical History

At eight months of age, the patient did not meet developmental milestones and was diagnosed with non-syndromic autism spectrum disorder. At approximately five years of age, the patient displayed an episode of repeated tripping and difficulty maintaining balance. A few days after, the patient displayed a convulsive seizure. The patient then underwent EEG monitoring which was classified as abnormal and the patient began treatment with valproate. No seizures were observed for three years, and valproate was tapered down at eight years of age. The patient then experienced a convulsive seizure, treatment with valproate was re-initiated, and seizures have not since been observed. At nine years of age, the patient underwent genetic testing in the form of whole exome sequencing which identified a *de novo* heterozygous variant in the *GRIN1* gene (RefSeq: NM_007327.3; c.1607A>G; p.Q536R) and hence a diagnosis of *GRIN1*-NDD. The center for Functional Evaluation of Rare Variants (CFERV) performed *in vitro* functional analysis on this variant and classified it as a loss-of-function variant, as described in the introduction.

### Generation and molecular characterization of *Grin1* ^Q536R/+^ mice

To evaluate the impact of the Q536R variant *in vivo, Grin1*^Q536R/+^ mice were created by CRISPR- Cas9 endonuclease-mediated transgenesis on a C57BL/6J genetic background (**Figure 1A**). The Gln536 residue is located in the S1 segment of the agonist binding domain (ABD), which harbours the binding site for glycine and is a conserved residue across vertebrate species, suggesting a critical role in NMDAR function (**Figure 1B, C**). *Grin1*^Q536R/+^ mice were born in expected Mendelian ratios and displayed normal survival and growth from PND 6-10 **(data not shown)**.

**Figure 1.**
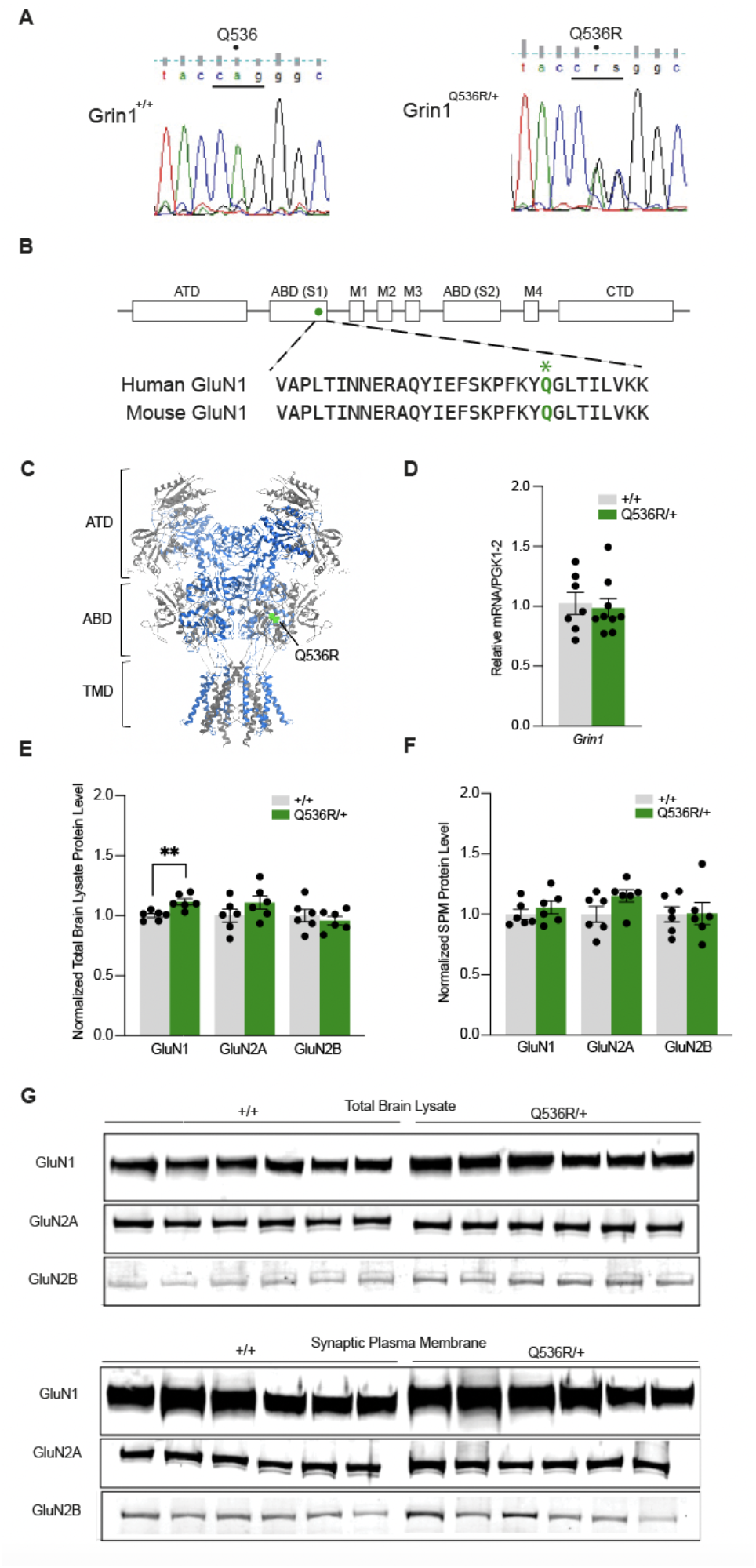
Substitution and molecular impacts of the *Grin1* Q536R variant in a mouse model. (**A**) Sanger sequencing of the A > G and G > C nucleotide mutations, resulting in the substitution of glutamine for arginine in Q536R/+ mice. (**B)** Linear schematic representation of the GluN1 subunit, demonstrating evolutionary conservation of amino acids between mouse and human. The location of the mutation in the S1 agonist-binding domain (ABD) is denoted by a circle, and the altered glutamine (Q) residue is denoted by an asterisk. ATD = amino terminal domain; CTD = carboxy terminal domain; M1-4 = transmembrane domains; agonist- binding domain (ABD) S1 and S2. **(C)** The domain architecture of the GluN1(grey)/GluN2A(blue) NMDAR with the location of the Q536R variant highlighted in green within the S1 ABD (Wang et al. 2021) built from human GluN1/GluN2A crystallographic data (PDB: 7EOS).**(D)** Relative quantification of forebrain*Grin1* mRNA expression normalized to housekeeping gene PGK1 and wildtype control (+/+) (n=7-9; p = 0.4870).**(E)** Relative levels of GluN1 (p = 0.0031), GluN2A (p = 0.1939) and GluN2B (p = 0.5062) protein in total brain lysate normalized to total protein stain and wildtype control (+/+) (n =6).**(F)** Relative levels of GluN1 (p = 0.4200), GluN2A (p = 0.0975) and GluN2B (p = 0.9541) protein in synaptic plasma membrane (SPM) fractions normalized to total protein stain and wildtype control (+/+) (n = 6).**(G)** Western blot of total brain lysate (top) and synaptic plasma membrane (bottom) fractions, probed for GluN1, GluN2A and GluN2B (n = 6). Data are expressed as mean ± SEM. **p < 0.01.

To examine the impact of the Q536R variant on mRNA transcripts and protein levels of NMDAR subunit components, RT-qPCR and western blot analysis were performed.

*Grin1*^Q536R/+^ mice did not differ in levels of forebrain *Grin1* mRNA compared to wildtype (p = 0.4870; **Figure 1D**). However, *Grin1*^Q536R/+^ displayed a modest but significant increase in GluN1 protein levels in total brain lysate (p = 0.0031) but not in synaptic plasma membrane fractions (p = 0.4200; **Figure 1E-G**). Conversely, GluN2A and 2B protein levels remained unchanged in *Grin1*^Q536R/+^ mice relative to wildtype in both total brain lysate (GluN2A, p = 0.1939; GluN2B, p = 0.5062) and synaptic plasma membrane fractions (GluN2A, p = 0.0975; GluN2B, p = 0.9541; **Figure 1E-G**).

### Adult *Grin1*^Q536R/+^ mice display deficits in NMDAR-mediated transmission

Given the critical role of NMDARs in excitatory neurotransmission, whole-cell patch-clamp electrophysiology was employed to explore the impact of the Q536R/+ variant on electrophysiological responses in the cortex. In response to bath application of NMDA (20 µM, 30s), layer 5 pyramidal neurons in the medial prefrontal cortex generated significantly less inward current in *Grin1*^Q536R/+^ than wildtype slices (p < 0.0001; **Figure 2A, B**). Additionally, hippocampal CA1 field potential recordings were used to probe differences in synaptic transmission between genotypes (**Figure 2C**). Stepwise increases in stimulus current intensity resulted in similar increases in fibre volley (FV) amplitude in slices from *Grin1*^Q536R/+^ and wildtype mice, and the corresponding slopes of the I/O curves plotting FV vs. stimulus intensity did not significantly differ between genotypes (p = 0.4033; **Figure 2D**). Therefore, presynaptic axon activation appeared largely unaffected in *Grin1*^Q536R/+^ mice. Similarly, AMPAR-mediated synaptic transmission as measured by the slope of the I/O curve plotting the AMPAR-fEPSP slope as a function of FV amplitude (p = 0.2893; **Figure 2E**) and paired-pulse facilitation, which is inversely related to neurotransmitter release probability (p = 0.4942; **Figure 2F**), did not differ significantly between genotypes. In contrast, recordings performed in low Mg^2+^ conditions and in the presence of AMPAR blocker NBQX revealed that slices from *Grin1*^Q536R/+^ displayed a significant reduction in the NMDAR-fEPSP slope as a function of FV amplitude compared to wildtype slices (p = 0.0007; **Figure 2G**). Taken together, these results suggest that the Q536R/+ variant does not confer major alterations in presynaptic function or AMPAR-mediated synaptic transmission but causes a significant deficit in NMDAR-mediated synaptic transmission in the hippocampal CA3-CA1 Schaffer collateral pathway.

**Figure 2.**
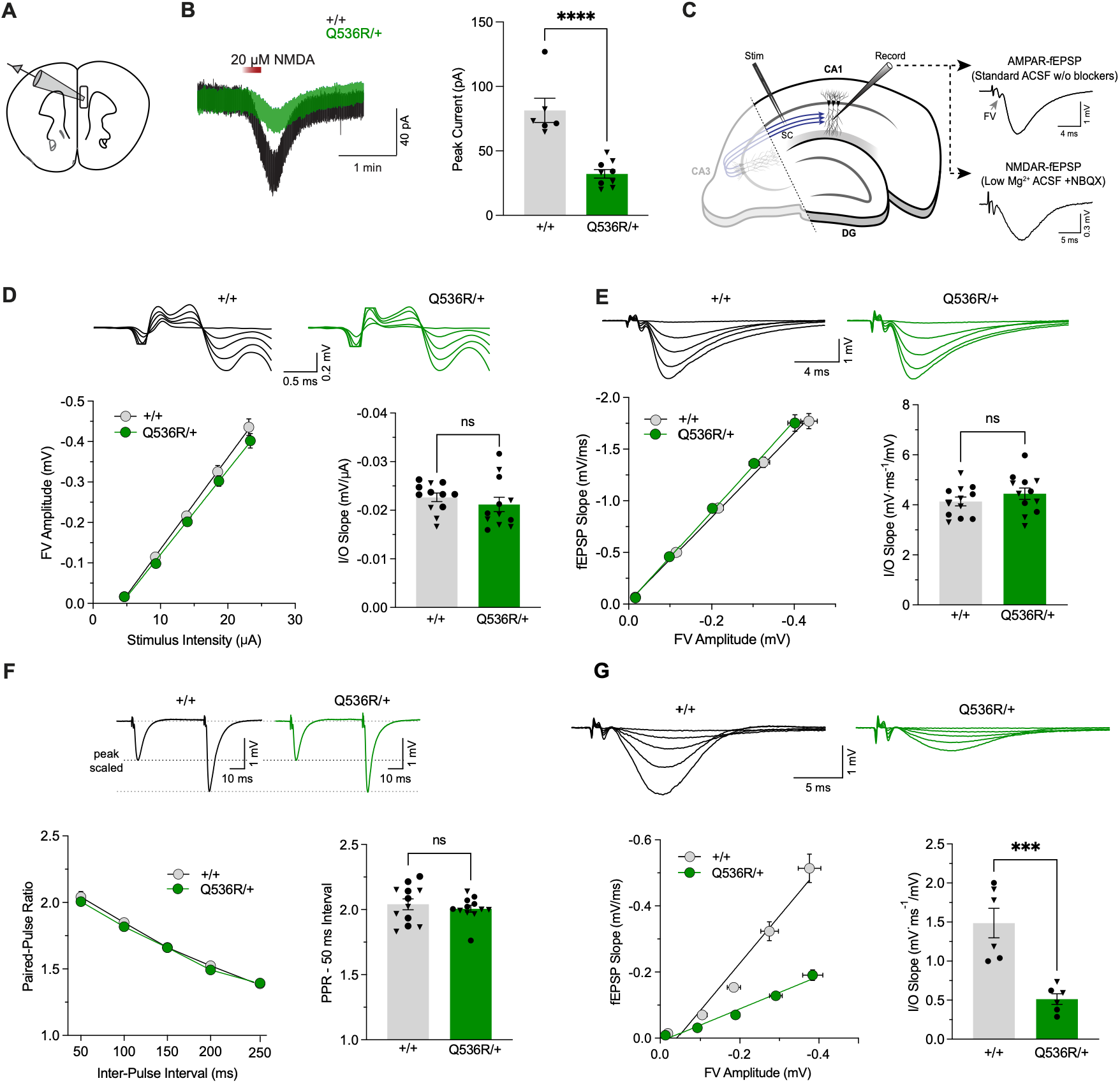
Deficits in NMDAR-mediated neurotransmission in the cortex and hippocampus of adult *Grin1* ^Q536R/+^ mice. **(A)** Schematic showing a coronal prefrontal cortex slice used to record NMDA stimulated currents in layer 5 pyramidal neurons. **(B)** Whole-cell currents evoked by bath application of NMDA (20 µM, 30s) to mPCF layer 5 pyramidal neurons was significantly reduced in Q536R/+ than +/+ slices (sexes combined: +/+ n = 6; Q536R/+ n = 9; p < 0.0001). **(C)** Schematic representation of a hippocampal slice displaying positions of the stimulating and recording electrodes for field potential recordings. CA3-CA1 fibre volleys (FV; indicated by grey arrow) and AMPA receptor-mediated fEPSPs (AMPAR-fEPSP) were recorded in standard ACSF (containing 1 mM Mg^2+^) without channel blockers. NMDA receptor-mediated fEPSPs (NMDAR-fEPSP) were recorded in low (0.1 mM) Mg^2+^ ACSF with NBQX (10 µM) to block AMPAR transmission. **(D)** Increasing stimulus intensity resulted in similar increases in FV amplitude in Q536R/+ and +/+ mice as shown in the input/output (I/O) curve (left). Genotypic differences were not observed in the corresponding linear regression fits (I/O slope) values, indicating largely intact presynaptic axon activation in Q536R/+ mice (right) (sexes combined: +/+ n = 12; Q536R/+ n = 12 [slices; 2 per animal]; p = 0.4033). Representative traces shown above. **(E)** I/O curves showing the relationship between FV amplitude and AMPAR-fEPSP slope in Q536R/+ and +/+ mice (left). Genotypic differences were not observed in the corresponding I/O slopes, indicating intact AMPAR-mediated synaptic transmission in Q536R/+ mice (right) (sexes combined: +/+ n = 12, Q536R/+ n = 12 [slices, 2 per animal]; p = 0.2893). Representative traces shown above. **(F)** Paired-pulse facilitation of AMPAR-fEPSPs measured across a range of inter-pulse intervals in Q536R/+ and +/+ mice (left). Genotypic differences were not observed in the paired pulse ratio (PPR, 50 ms inter-pulse interval) at any tested time points, suggesting no detectable differences in neurotransmitter release probability (right) (sexes combined: +/+ n = 12; Q536R/+ n =12 [slices, 2 per animal]; p = 0.4942). Representative traces shown above. **(G)** I/O curves showing the relationship between FV amplitude and NMDAR-fEPSP slope in Q536R/+ and +/+ mice (left). Q536R/+ mice displayed a significantly reduced I/O slope compared to +/+, indicative of deficient NMDAR-mediated synaptic transmission (right). (sexes combined: +/+ n = 6, Q536R/+ n = 6 [slices, 1 per animal]; p = 0.0007). Representative traces shown above. Data are expressed as mean ± SEM. ***p < 0.001; ****p < 0.0001; ns: not significant. Males and females are represented by circle and triangle symbols, respectively.

### Adult *Grin1*^Q536R/+^ mice display deficits in long-term potentiation but have intact depotentiation

To assess the impact of the Q536R/+ variant on synaptic plasticity mechanisms, long-term potentiation (LTP) was induced using a compressed theta burst stimulation (cTBS) protocol at CA3-CA1 synapses. *Grin1*^Q536R/+^ mice had a significantly reduced magnitude of LTP (average fEPSP slope change 80-90 min post TBS) relative to wildtype mice (p = 0.0027; **Figure 3A**).

**Figure 3.**
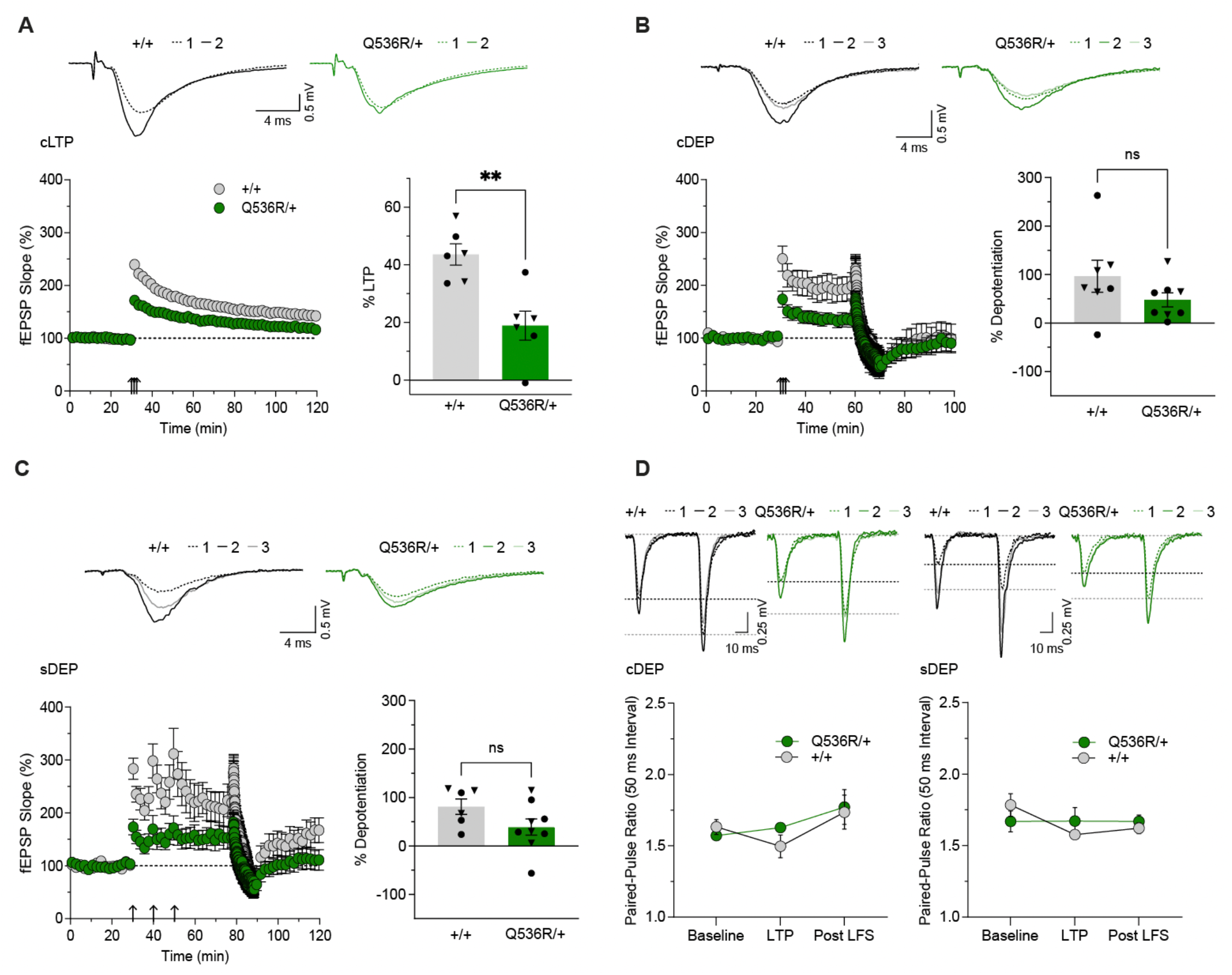
Adult *Grin1* ^Q536R/+^ mice display deficits in long-term potentiation but intact depotentiation in the hippocampus. **(A)** Time-course plot showing the change in AMPAR-fEPSP slope (plotted as % of baseline) following delivery of compressed theta burst stimulation (cTBS) in Q536R/+ and +/+ slices (left). Summarized magnitude of LTP (% change from baseline) measured 80 – 90 min post cTBS (right). LTP was significantly reduced in Q536R/+ mice (sexes combined: +/+ n = 6; Q536R/+ n = 6 [slices, 1 per animal]; p = 0.0027). Representative traces recorded before (1) and after (2) TBS are shown above. **(B)** Time-course plot showing the change in AMPAR-fEPSP slope (plotted as % of baseline) following cTBS administration and subsequent DEP induction (cLTP-DEP) in Q536R/+ and +/+ slices (left). Summarized magnitude of the percent depotentiation (right) (sexes combined: +/+ n = 7, Q536R/+ n = 8 [slices, 1 per animal]; p = 0.1806). Representative traces recorded for baseline (1), LTP (2) and depotentiation (3) are shown above. **(C)** Time-course plot showing the change in AMPAR-fEPSP slope (plotted as % of baseline) following spaced TBS (sTBS) and subsequent DEP induction (sLTP-DEP) in Q536R/+ and +/+ slices (left). Summarized magnitude of percent depotentiation (right) (sexes combined: +/+ n = 6, Q536/+ n = 9 mice [slices, 1 per animal]; p = 0.1050). **(D)** Paired-pulse facilitation of AMPAR-fEPSPs measured at baseline, after LTP induction, and post LFS administration for the cLTP-DEP (left) and sLTP-DEP (right) recordings in Q536R/+ and +/+ mice. Genotypic differences were not observed in the paired pulse ratio (PPR, 50 ms inter-pulse interval) at any time point tested (two-way RM-ANOVA: time × genotype interaction: F[2,16] = 0.6953, p = 0.5134 [cLTP-DEP]; F[2,14] = 1.606, p = 0.2355 [sLTP-DEP]). (sexes combined: cLTP-DEP: +/+ n = 5, Q536R/+ n = 5 [slices, 1 per animal]; sLTP-DEP: +/+ n = 3, Q536R/+ n = 6 [slices, 1 per animal]). Data are expressed as mean ± SEM. **p < 0.01; ns: not significant. Males and females are represented by circle and triangle symbols, respectively.

These data are consistent with the previously observed impairments in NMDAR-mediated synaptic transmission.

Next, to test whether synaptic weakening is similarly impaired in adult *Grin1*^Q536R/+^ mice, we assessed LTP reversal, or synaptic depotentiation (DEP). Like LTP, hippocampal DEP triggered by low-frequency stimulation (LFS) is NMDAR-dependent and can be readily induced in adult animals (Huang et al., 2001; Zhang et al., 2009; Ge et al., 2019). In wildtype mice, LTP induced by cTBS (cLTP) was depotentiated by a 2Hz LFS protocol delivered 30 minutes after induction to approximately baseline levels (**Figure 3B**), consistent with previous reports (Park et al., 2019). Interestingly, despite the reduced magnitude of cLTP, cLTP-DEP also remained intact in *Grin1*^Q536R/+^ mice (**Figure 3B**), with no statistically significant differences detected in the percent DEP between genotypes (p = 0.1806).

We then tested whether the Q536R/+ variant may specifically impair DEP of LTP induced by a spaced TBS (sTBS) protocol, since DEP is sensitive to the type of LTP induced (Woo and Nguyen, 2003; Pauli et al., 2026). Indeed, it has been previously shown that LTP induced with sTBS (sLTP) is more resistant to depotentiation (sLTP-DEP) (Park et al., 2019). Consistent with these previous studies, we observed that, in wildtype mice, sLTP is resistant to DEP and responses remained elevated above baseline levels following the 2 Hz LFS depotentiation protocol (**Figure 3C**). In *Grin1*^Q536R/+^ mice, despite reduced sLTP, sLTP-DEP remained intact (p = 0.1050 vs. wildtype sLTP-DEP; **Figure 3C**). To summarize, *Grin1*^Q536R/+^ mice display impaired LTP with both induction protocols (compressed and spaced) but the process of DEP is intact in these mutant mice.

Lastly, to assess possible differences in presynaptic alterations during plasticity induction, we monitored the paired-pulse ratios (PPRs) of fEPSPs evoked 50 ms apart at baseline, after LTP induction, and after DEP induction. No statistically significant genotype differences were observed in the PPR at any measured timepoint (two-way RM-ANOVA: time × genotype interaction: F[2,16] = 0.6953, p = 0.5134 [cLTP-DEP]; F[2,14] = 1.606, p = 0.2355 [sLTP-DEP]; **Figure 3D**). Therefore, the observed findings are unlikely to be due to presynaptic alterations. Together, these findings demonstrate that although LTP is reduced in *Grin1*^Q536R/+^ mice, DEP remains intact regardless of the type of LTP induced.

### Alterations to dentate gyrus morphology are present in adult *Grin1*^Q536R/+^ mice

NMDARs are highly expressed in the cerebral cortex and hippocampus subfields, and related neuroanatomical abnormalities have been observed in some individuals with *GRIN1*-NDD (Fry et al., 2018; Platzer and Lemke, 2019). Thus, the potential for gross histological changes in these regions was examined in *Grin1*^Q536R/+^ mice. Compared to wildtype mice, no significant differences were observed in cortical thickness (p = 0.6620), pyramidal cell layer area (p = 0.1472), thickness of the hippocampus proper (p = 0.9880), or thickness of the CA1 (p = 0.5760) and CA2 (p = 0.2216) subfields (**Figure 4A-E**). However, *Grin1*^Q536R/+^ mice displayed a significant decrease in the granule cell layer area of the dentate gyrus (p = 0.0048), and a corresponding trend towards a decreased thickness of the same region (p = 0.0871) (**Figure 4B,D,E**).

**Figure 4.**
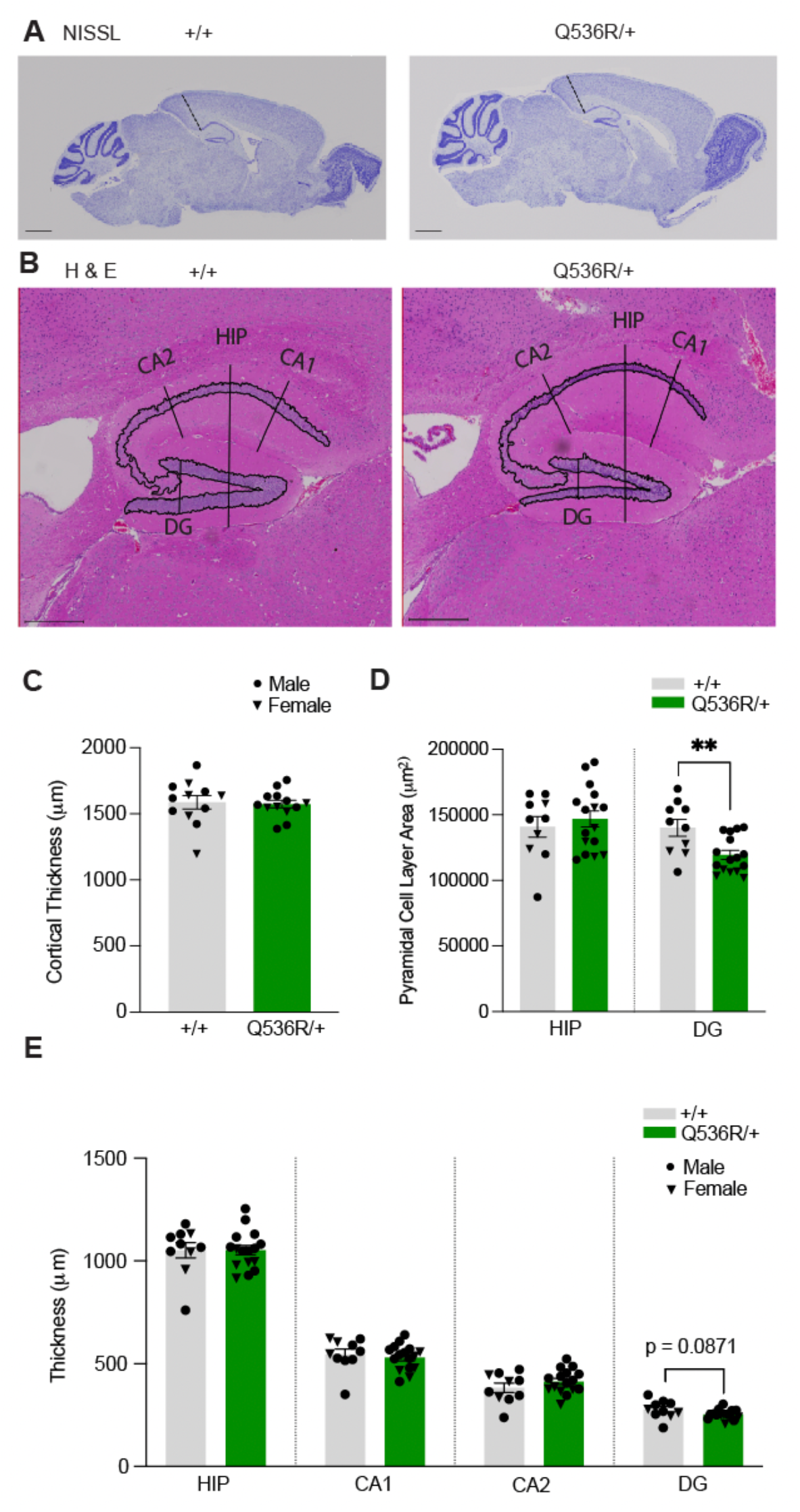
Alterations to dentate gyrus morphology in adult *Grin1* ^Q536R/+^ mice. **(A)** Representative images of Nissl-stained sagittal brain sections of +/+ and Q536R/+ mice with cortical thickness measurements indicated by the dashed lines (scale bars = 1 mm).**(B)** Representative images of H&E-stained hippocampus proper, CA1, CA2 and dentate gyrus regions in +/+ and Q536R/+ mice with thickness and cell layer area measurements indicated by solid lines and shaded areas, respectively (scale bars = 400 μm).**(C)** Cortical thickness was not significantly different between genotypes (sexes combined: +/+ n = 12, Q536R/+ n = 13; p = 0.6620).**(D)** Q536R/+ mice displayed a significantly reduced area of the dentate gyrus granule cell layer (sexes combined: +/+ n = 10; Q536R/+ n = 16; p = 0.0048), but not of the hippocampus proper pyramidal cell layer (p = 0.9880).**(E)** Thickness of the dentate gyrus trended towards a reduction in Q536R/+ compared to +/+ mice (sexes combined: +/+ n = 10; Q536R/+ n = 16; p = 0.0871). Data are expressed as mean ± SEM. **p < 0.01. Males and females are represented by circle and triangle symbols, respectively.

### Grin1^Q536R/+^ mice display altered vocal communication in the early post-natal period

At PND 6, *Grin1*^Q536R/+^ mice displayed intact righting reflexes (20 ± 4.7 seconds) that did not significantly differ from wildtype mice (21 ± 3.6 seconds) (p = 0.8911; **Figure 5A**). However, in response to maternal isolation, *Grin1*^Q536R/+^ mice emitted significantly less ultrasonic vocalizations compared to wildtype littermates (two-way ANOVA: main effect of genotype: F[1,61] = 5.692; p = 0.0202) (**Figure 5B**). At PND 21, *Grin1*^Q536R/+^ mice did not display significant changes in bodyweight (p = 0.8182; **Figure S1A**) or changes in muscle tone as indicated by holding impulse in the wire hang task (p = 0.5239; **Figure 5C**).

**Figure 5.**
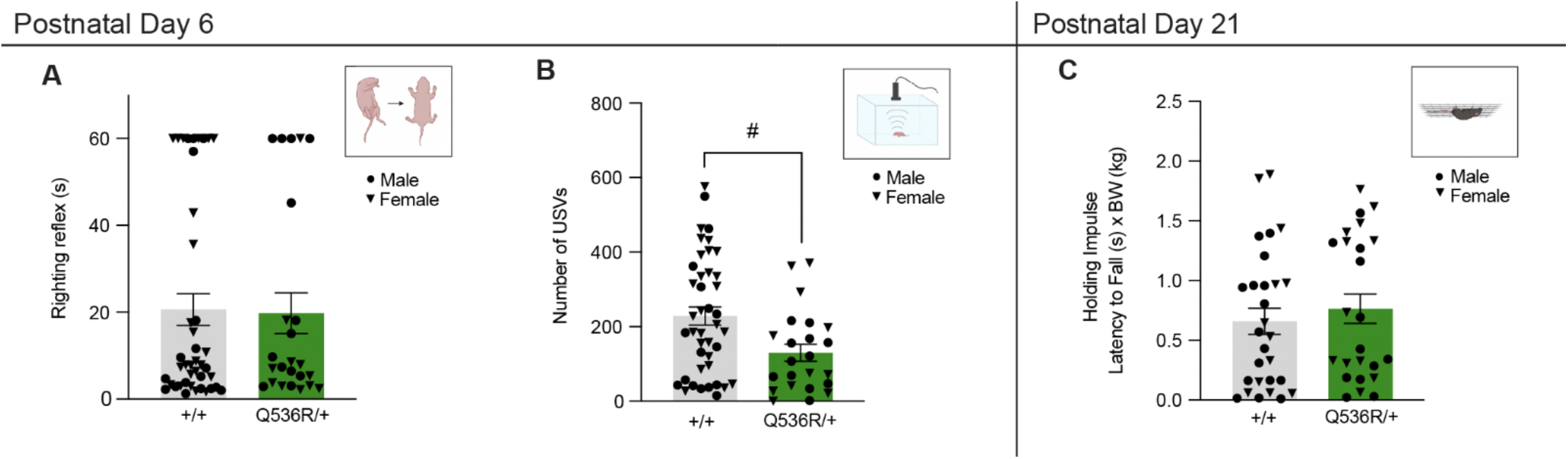
*Grin1* ^Q536R/+^ mice display alterations in vocal communication in response to maternal isolation. **(A)** At PND 6, Q536R/+ mice displayed righting reflexes similar to +/+ mice (+/+ n = 42, M = 16, F = 26; Q536R/+ n = 24, M = 13, F = 11; p = 0.8911).**(B)** Q536R/+ mice displayed a reduction in the number of ultrasonic vocalizations elicited in response to maternal isolation compared to +/+ mice at PND 6 (+/+ n = 42, M = 16, F = 26; Q536R/+ n = 24, M =13, F = 11; two-way ANOVA: main effect of genotype: F[1,61] = 5.692; p = 0.0202).**(C)** At PND 21, holding impulse in the wire hang task was not significantly different between genotypes (+/+ n = 29, M = 17, F = 12; Q536R/+ n = 24, M = 12, F= 12; p = 0.5239). Data are expressed as mean ± SEM. Main effect # p < 0.05. Males and females are represented by circle and triangle symbols, respectively.

### Adult *Grin1*^Q536R/+^ mice display modest and sex specific behavioural phenotypes

In adulthood, *Grin1*^Q536R/+^ mice did not display alterations in bodyweight compared to wildtype mice (two-way ANOVA: main effect of genotype: F[1,55] = 0.5793, p = 0.4498; **Figure S1B**). However, male *Grin1*^Q536R/+^ mice displayed a significant deficit in the holding impulse measured in the wire hang task (two-way ANOVA: genotype x sex interaction: F[1,55] = 4.467, p = 0.0391; Tukey’s post hoc test: males: p = 0.0470) (**Figure 6A**). In the open field test, the total distance travelled did not differ between genotypes (two-way ANOVA: main effect of genotype: F[1,44] = 0.03347, p = 0.8557), but male *Grin1*^Q536R/+^ mice travelled a significantly increased distance compared to *Grin1*^Q536R/+^ females (two-way ANOVA: genotype × sex interaction: F[1,44] = 6.237, p = 0.0163; Tukey’s post hoc test: *Grin1*^Q536R/+^ males vs females: p = 0.0124; **Figure 6B,C**).

**Figure 6.**
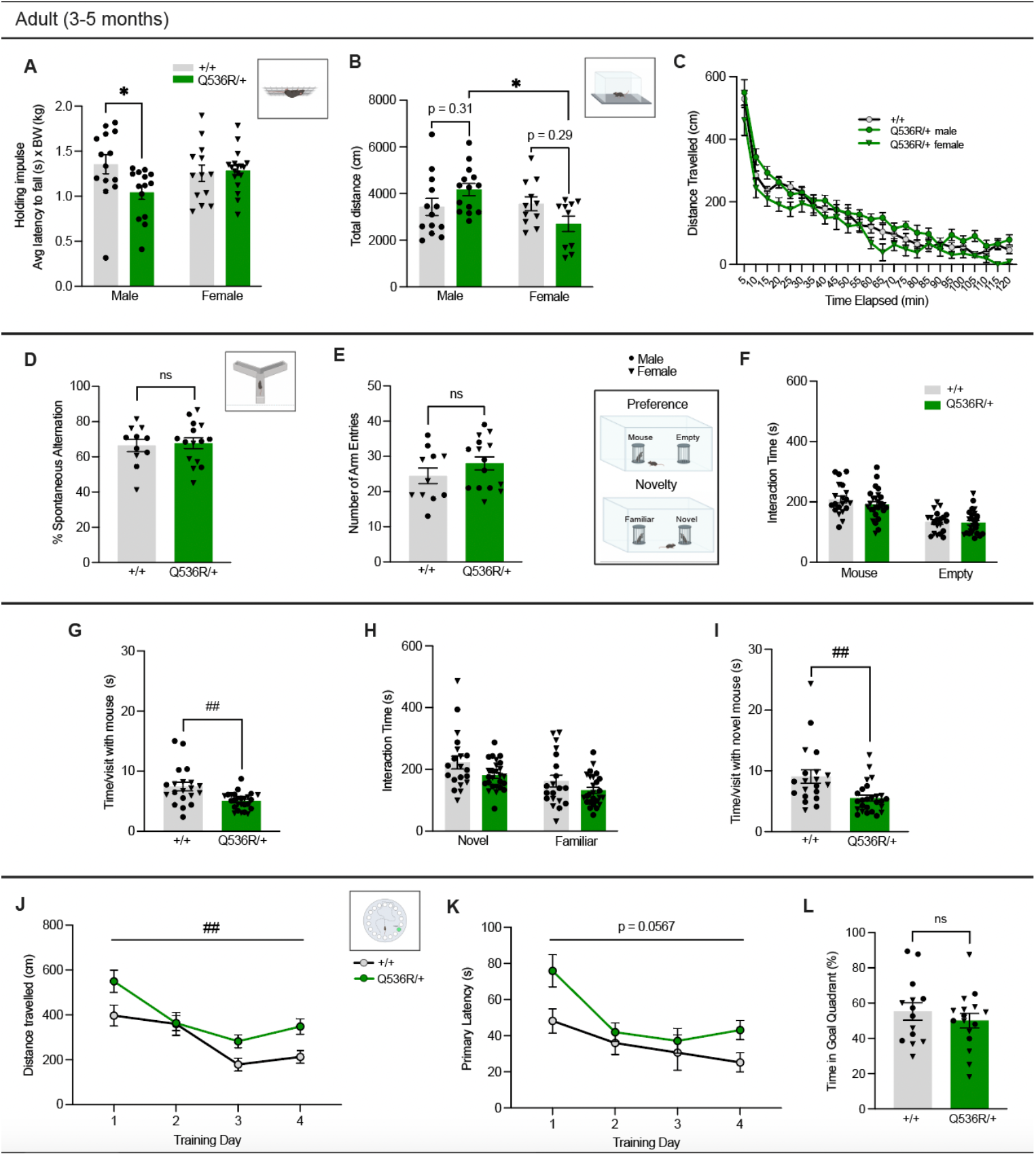
Adult *Grin1* ^Q536R/+^ mice display sex specific phenotypes. **(A)** Male but not female Q536R/+ mice, displayed significantly reduced holding impulse in the wire hang task relative to +/+ (+/+ n = 27, M = 14, F = 13; Q536R/+ n = 32, M = 14, F = 18; two-way ANOVA: genotype × sex interaction: F[1,55] = 4.467, p = 0.0391; Tukey’s post hoc tests: male +/+ vs. male Q536R/+, p = 0.0470). **(B)** In the open field test, distance travelled over 2-hours was not significantly different between genotypes (+/+ n = 24, M = 13, F = 11; Q536R/+ n = 24, M = 14, F = 10; two-way ANOVA: main effect of genotype: F[1,44] = 0.03347, p = 0.8557), but male Q536R/+ mice travelled significantly more distance compared to Q536R/+ females (two-way ANOVA: main effect of sex: F[1,44] = 4.329, p = 0.0433; significant genotype x sex interaction: F[1,44] = 6.237, p = 0.0163; Tukey’s post hoc tests: male Q536R/+ vs. female Q536R/+, p = 0.0124). **(C)** Distance travelled over time during the 2-hour open field test (+/+ n = 24, M = 13, F = 11; Q536R/+ n = 24, M = 14, F = 10). **(D,E)** In the Y-maze test of working memory, Q536R/+ mice did not display significant differences in % spontaneous alternation (+/+ n = 11, M = 6, F = 5; Q536R/+ n = 15, M = 8, F = 7; two-way ANOVA: main effect of genotype: F[1,22] = 0.04462, p = 0.8346), or the number of arm entries entered during the duration of the test compared to +/+ mice (two-way ANOVA: main effect of genotype: F[1,22] = 1.395, p = 0.2501). **(F)** In the social interaction test, Q536R/+ mice had intact preference for social interaction, spending significantly more time with the mouse than the empty cup, as was seen in +/+ mice (+/+ n =20, M = 11, F = 9; Q536R/+ n = 26, M = 15, F = 11; three-way ANOVA: main effect of presence of mouse: F[1,42] = 33.65, p < 0.0001). **(G)** However, Q536R/+ spent a significantly reduced amount of time with the mouse per visit (two-way ANOVA: main effect of genotype: F[1,42] = 11.52, p = 0.0015). **(H)** Similarly, Q536R/+ mice had an intact preference for social novelty, spending significantly more time with the novel than the familiar mouse, as was seen in +/+ mice (+/+ n = 20, M = 11, F = 9; Q536R/+ n = 26, M = 15, F = 11; three-way ANOVA: main effect of novelty of mouse: F[1,42] = 8.243, p = 0.0064; main effect of genotype: F[1,42] = 13.03, p = 0.0008). **(I)** However, Q536R/+ spent a significantly reduced amount of time with the novel mouse per visit (+/+ n = 20, M = 11, F = 9; Q536R/+ n = 26, M = 15, F = 11; two-way ANOVA: main effect of genotype: F[1,42] = 10.06, p = 0.0028). **(J)** In the Barnes Maze, Q536R/+ mice travelled significantly increased distance to reach the goal box during acquisition trials (+/+ n = 15, M = 8, F = 6; Q536R/+ n = 17, M = 5, F = 11; two-way RM-ANOVA: main effect of genotype: F[1,28] = 8.358, p = 0.0073, main effect of training day: F[3,84] = 18.84, p < 0.0001). **(K)** Similarly, Q536R/+ trended toward increased latency to reach the goal box during acquisition trials (+/+ n = 14, M = 8, F = 6; Q536R/+ n = 16, M = 5, F = 11; two-way RM-ANOVA: trend toward main effect of genotype: F[1,28] = 3.952, p = 0.0567, main effect of training day: F[3,84] = 12.18, p < 0.0001). **(L)** However, Q536R/+ mice spent a similar % of time spent in the goal quadrant compared to +/+ mice on the probe trial (unpaired t test, p=0.4222). Data are expressed as mean ± SEM. *p < 0.05. Main effect ## p < 0.01. ns: not significant. Males and females are represented by circle and triangle symbols, respectively.

Genotype differences in spontaneous alternation behaviour were not observed in the Y-maze (two-way ANOVA: main effect of genotype: F[1,22] = 0.04462, p = 0.8346), indicating intact working memory in adult *Grin1*^Q536R/+^ mice (**Figure 6D**). Additionally, *Grin1*^Q536R/+^ and wildtype mice displayed similar levels of exploration and did not differ in the number of arm entries made during the duration of the Y-maze test (two-way ANOVA: main effect of genotype: F[1,22] = 1.395, p = 0.2501; **Figure 6E**).

The social interaction test was used to investigate the innate preference of mice to investigate a stranger mouse of the same sex rather than an empty cage (social preference) and for investigating a novel rather than familiar conspecific (social novelty preference). *Grin1*^Q536R/+^ mice displayed an intact social preference and spent significantly more time exploring the mouse than the empty enclosure (three-way ANOVA: main effect of presence of mouse: F[1,42] = 33.65, p < 0.0001; **Figure 6F**). However, the quality of social interaction engaged in by *Grin1*^Q536R/+^ mice appeared to be diminished, as demonstrated by a significant reduction in the amount of time spent with the same-sex mouse on each visit compared to wildtype (two-way ANOVA: main effect of genotype: F[1,42] = 11.52, p = 0.0015; **Figure 6G**). *Grin1*^Q536R/+^ mice displayed social novelty preference, spending more time with the novel mouse compared to the familiar mouse (three-way ANOVA: main effect of novelty of mouse: F[1,42] = 8.243, p = 0.0064; **Figure 6H**). However, there was a significant main effect of genotype in this task, indicating *Grin1*^Q536R/+^ mice had reduced interaction time overall compared to wildtype mice, regardless of whether the mouse was novel or familiar (main effect of genotype: F[1,42] = 13.03, p = 0.0008; **Figure 6H**). This could reflect reduced social interest for both the novel and familiar mice leading to less meaningful interaction with test mice as compared to wildtype. Accordingly, *Grin1*^Q536R/+^ mice spent significantly less time with the novel mouse on each visit as compared to wildtype mice (two-way ANOVA: main effect of genotype: F[1,42] = 10.06, p = 0.0028; **Figure 6I**).

Next, *Grin1*^Q536R/+^ mice were tested in the Barnes maze, a spatial learning and memory task that requires mice to learn the position of a goal box to escape an open and brightly lit test arena. During training, *Grin1*^Q536R/+^ mice travelled a significantly greater distance to reach the goal box (two-way RM-ANOVA: main effect of genotype: F[1,28] = 8.358, p = 0.0073, main effect of training day: F[3,84] = 18.84, p < 0.0001; **Figure 6J**). This was mirrored by a notable pattern of increase in latency to reach the goal box during acquisition trials (two-way RM-ANOVA: trend toward main effect of genotype: F[1,28] = 3.952, p = 0.0567, main effect of training day: F[3,84] = 12.18, p < 0.0001; **Figure 6K**). Despite this, on the probe trial, *Grin1*^Q536R/+^ and wildtype mice spent a similar percentage of time in the goal quadrant (p = 0.4222; **Figure 6L**). Thus, *Grin1*^Q536R/+^ mice learn the spatial location of the goal box at a slower rate but ultimately form a spatial memory of it on par with wildtype mice.

Lastly, the presence of behavioural convulsions was evaluated in *Grin1^Q536R^*^/+^ mice through once-a-week handling. During weekly handling by experimenters for a duration of 12 weeks, spontaneous convulsions were absent in wildtype mice and were occasionally elicited in *Grin1*^Q536R/+^ mice. In week 4 of monitoring, where seizures were the most prominent, only 21% (3/14) of *Grin1*^Q536R/+^ mice displayed seizures **(Figure S2A).** Interestingly, one male *Grin1*^Q536R/+^ mouse reliably seized upon handling and displayed seizures of advanced severity (up to category 6) **(Figure S2B,C; Table S1)**.

### Older adult *Grin1* ^Q536R/+^ mice of both sexes display hyperactivity

*Grin1*^Q536R/+^ mice at 9-13 months of age (older adult) did not display differences in bodyweight relative to wildtype mice (two-way ANOVA: main effect of genotype: F[1,39] = 3.687, p = 0.0622; **Figure S1C)**. However, we found that sexually dimorphic locomotor phenotypes observed in adult *Grin1*^Q536R/+^ mice dissipate with age, older *Grin1*^Q536R/+^ of both sexes travelled a significantly greater total distance in the open field test compared to wildtype (two-way ANOVA: main effect of genotype: F[1,39] = 10.66, p = 0.0023) (**Figure 7A**). There was a statistically significant interaction between the effects of genotype and time on distance travelled in the open field test (two-way RM-ANOVA; genotype × time interaction: F[23,943] = 2.813, p < 0.0001), with aged *Grin1*^Q536R/+^ mice travelling significantly greater distance than wildtype mice within each 5-minute time bin from 5-20 min, 35, and 85 min (Tukey’s post hoc tests: 5, p < 0.0001; 10, p < 0.0001; 15, p < 0.0001; 20, p = 0.0060; 35, p = 0.0024 85, p = 0.0358; **Figure 7B**).

**Figure 7.**
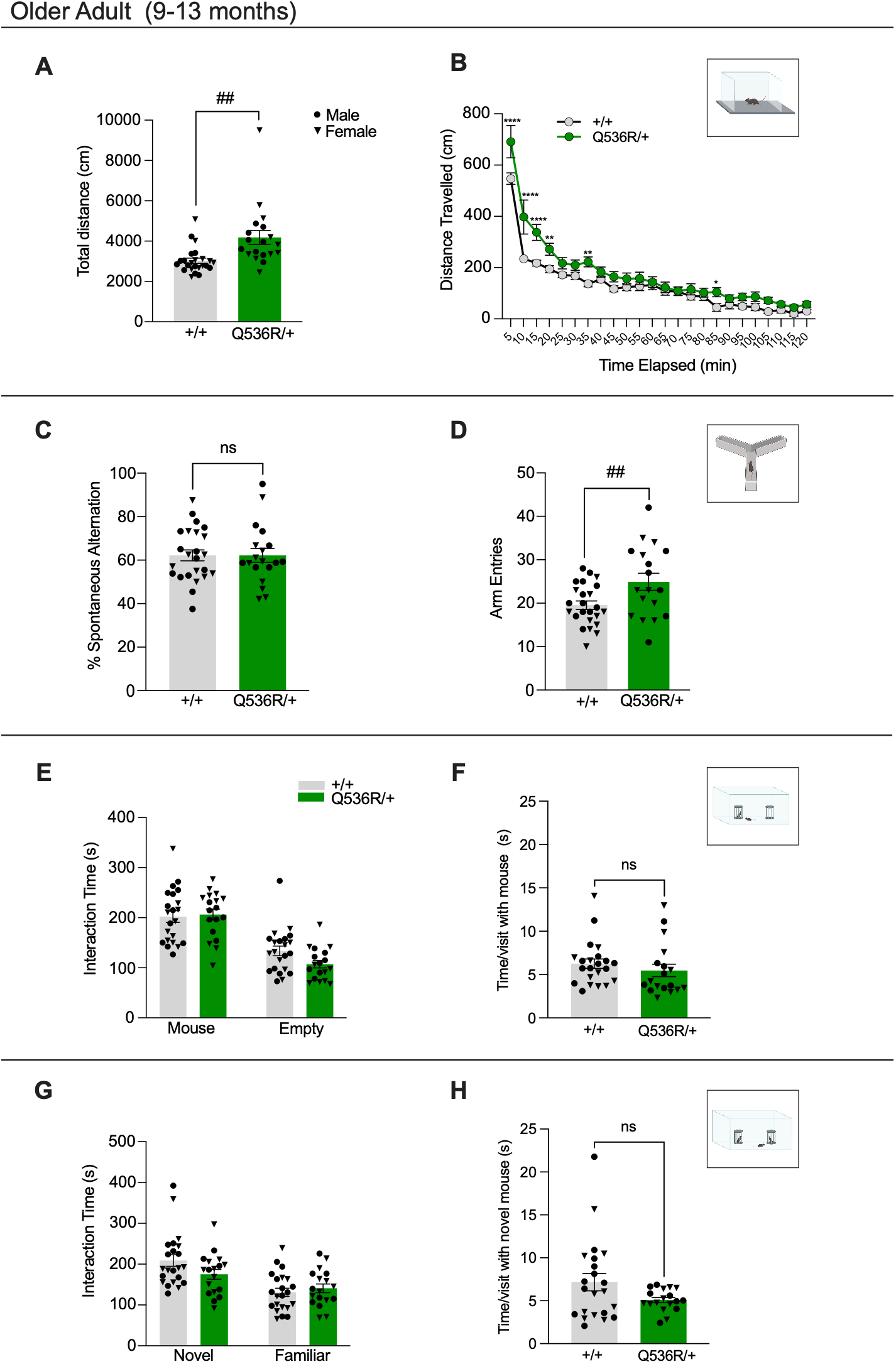
Older adult *Grin1* ^Q536R/+^ mice display hyperactivity. **(A)** In the open field test, older adult Q536R/+ mice of both sexes travelled significantly greater distance than +/+ mice (+/+ n = 24, M = 13, F = 11; Q536R/+ n = 19, M = 9, F = 10; two-way ANOVA main effect of genotype: F[1,39] = 10.66, p = 0.0023). **(B)** Distance traveled over time during the 2 hour open field test, with Q536R/+ mice travelling significantly greater distance than +/+ mice in each 5-min bin from 5-20 min, 35, and 85 min (sexes combined: +/+ n = 24, Q536R/+ n = 19; two-way RM-ANOVA genotype × time interaction: F[23,943] = 2.813, p < 0.0001; Tukey’s post hoc test: 5, p < 0.0001; 10, p < 0.0001; 15, p < 0.0001; 20, p = 0.0060; 35, p = 0.0024; 85, p = 0.0358; main effect of time: F[23, 943] = 100.5, p < 0.0001; main effect of genotype: F[1,41] = 11.37, p = 0.0016). **(C)** In the Y-maze, percentage of spontaneous alternation did not differ between genotypes (+/+ n = 24, M = 13, F = 11; Q536R/+ n = 19, M = 9, F = 10; two-way ANOVA main effect of genotype: F[1,39] = 0.001413, p = 0.9702). **(D)** However, older adult Q536R/+ mice of both sexes displayed a significantly increased number of arm entries compared to +/+ mice (two-way ANOVA main effect of genotype: F[1,38] = 8.159, p = 0.0069). **(E)** In the social interaction test, older adult Q536R/+ mice had an intact preference for social interaction (+/+ n = 22, M = 12, F = 10; Q536R/+ n = 18, M = 8, F = 10; three-way ANOVA: main effect of presence of mouse: F[1,36] = 44.70, p < 0.0001) and **(F)** spent a similar amount of time with the mouse per visit compared to +/+. **(G)** Similarly, older adult Q536R/+ mice had an intact preference for social novelty, spending significantly more time with the novel than the familiar mouse, as was seen in +/+ mice (three-way ANOVA: main effect of novelty of mouse: F[1,36] = 15.73, p = 0.0003) and (**H**) spent a similar amount of time with the novel mouse per visit. Data are expressed as mean ± SEM. Main effect: ## p < 0.01. Males and females are represented by circle and triangle symbols, respectively.

Moreover, as was observed in early adulthood, older *Grin1*^Q536R/+^ mice did not display genotype differences in terms of the percentage of spontaneous alternations in the Y-maze task (two-way ANOVA: main effect of genotype: F[1,39] = 0.001413, p = 0.9702). However, aging did alter the exploration pattern of *Grin1*^Q536R/+^ mice, which exhibited a significantly increased number of arm entries during the task relative to wildtype mice (two-way ANOVA: main effect of genotype: F[1,38] = 8.159, p = 0.0069; **Figure 7C,D**).

Lastly, older adult *Grin1*^Q536R/+^ mice continued to display social preference (three-way ANOVA: main effect of presence of mouse: F[1,36] = 44.70, p < 0.0001; (**Figure 7E**) and social novelty preference (three-way ANOVA: main effect of novelty of mouse: F[1,36] = 15.73, p = 0.0003; **Figure 7G**). However, the quality of social interaction appeared to improve with age, as genotype differences were no longer observed in the amount of time spent on each visit with the probe mouse during the social preference task (**Figure 7F**) or the novel mouse during the social novelty preference task (**Figure 7H**).

## Discussion

Here, we have reported the generation and characterization of a mouse model carrying the heterozygous Q536R variant of *Grin1* (*GRIN1*) found in a male patient with DD/ASD. We assessed the pathogenicity of the heterozygous Q536R variant at molecular, synaptic, histological and behavioural levels. Besides a modest increase of GluN1 expression in brain tissue lysate, the mRNA and protein analyses showed no other changes in the expression of major NMDAR subunits (GluN1, GluN2A, and GluN2B) in either brain tissue lysate or synaptic plasma membrane fraction. These results collectively suggest normal receptor trafficking and subcellular expression in the presence of the Q536R/+ variant. However, impaired NMDAR function was observed as shown by reduction of NMDA-induced currents in the prefrontal cortex layer 5 pyramidal cells, as well as decreased NMDAR-mediated synaptic transmission and reduced LTP at the hippocampal Schaffer collateral-CA1 synapses. Since NMDAR subunit expression was normal, the functional impairment of neuronal NMDARs is likely a dominant negative LoF effect due solely to the disruption of glutamate (-3.7-fold) and glycine (-3432-fold) potency as previously described (CFERV, Emory University, 2019).

The *Grin1*^Q536R/+^ mice afforded an opportunity to dissect the contribution of glycine binding to aspects of synaptic strengthening and weakening as studied by electrical stimulation protocols in hippocampal slice preparations. Although *Grin1*^Q536R/+^ mice demonstrated significantly reduced hippocampal LTP, DEP remained intact. Thus, *Grin1*^Q536R/+^ mice exhibit altered synaptic plasticity in which synaptic strengthening is impaired, but synaptic weakening still functions. Since the primary deficit of NMDARs imparted by the Q536R variant in these mice is the absence of glycine binding, our findings support a requirement for concurrent glutamate and glycine binding during LTP induction, yet may reveal that only glutamate binding is necessary for DEP. Indeed, NMDAR signaling independent of ion flux (non-ionotropic NMDAR signaling) can be induced upon glutamate binding in the presence of glycine site-specific antagonists, and it is specifically involved in hippocampal long-term depression and spine shrinkage (Nabavi et al., 2013; Stein et al., 2015; Wong and Gray, 2018). Non-ionotropic NMDAR signaling was also shown to play a role in synaptic DEP induced at Schaffer collateral- CA1 synapses (Latif-Hernandez et al., 2016; Pauli et al., 2026). *Grin1*^Q536R/+^ mice would therefore be expected to have intact non-ionotropic NMDAR signaling associated with long-term depression and DEP but would have impaired ionotropic signaling necessary for LTP induction. Importantly, this could explain why *Grin1*^Q536R/+^ mice displayed slower learning yet normal spatial memory acquisition, as hippocampal DEP has been correlated with novelty exploration and spatial-based learning (Xu et al., 1998; Straube et al., 2003; Qi et al., 2013; Ge et al., 2019; Pauli and Bonin, 2026).

The *Grin1*^Q536R/+^ mouse model also allowed us to investigate neuroanatomical changes that have been observed with NMDAR manipulations and in *GRIN1* patients. Polymicrogyria (PMG), a malformation of cortical development, has been reported in a subset of *GRIN1* patients (Fry et al., 2018; Brock et al., 2023). However, we observed no changes in cortical thickness or organization, predicting a reduced likelihood of cortical malformation associated with the Q536R/+ variant. Indeed, PMG-associated *GRIN1* variants were reported to cluster in the S2, M3, and S1-M1 linker regions (Fry et al., 2018), whereas the Q536R/+ variant resides in the S1 domain. To date, no *GRIN1* S1 domain variants have been directly linked to polymicrogyria. While cortical anatomy was normal, *Grin1*^Q536R/+^ mice did show specific reductions in the cell body area of the dentate gyrus region of the hippocampus, a region associated with adult neurogenesis. A similar phenotype has been observed in mice with region-specific knockout of the *Grin1* gene in granule cells of the dentate gyrus and was attributed to observed impairments in postnatal and adult neurogenesis (Åmellem et al., 2021). Further investigation of neurogenesis is required to determine the *in vivo* pathological mechanisms conferred by the Q536R/+ variant.

We lack a comprehensive understanding of the natural history of *GRIN1*-NDD, but mouse models can help inform the impact of NMDAR dysfunction over time. To understand the progressive consequences of the Q536R/+ variant, we monitored the body weight and conducted behavioral analysis across the lifespan of *Grin1*^Q536R/+^ mice. We focused on behavioral domains reflecting the patient’s clinical manifestations: muscle strength, cognition, and social interactions. At PND 21, indications of hypotonia and developmental delay in *Grin1*^Q536R/+^ mice were not evident. However, in adulthood, a sex-specific deficit in the wire hang task became evident in male *Grin1*^Q536R/+^ mice, reflecting the hypotonia observed in the male patient. A sex- specific difference was also observed in the open field test, where male *Grin1*^Q536R/+^ mice were significantly more hyperactive than female *Grin1*^Q536R/+^ mice. Similarly, sex-specific alterations have also been observed in *Grin1*KD mice (a mouse model with 90% GluN1 reduction) and the *Grin2b*^L825V/+^ patient variant model (Milenkovic et al., 2014; Serra et al., 2024). These findings underscore the importance of examining sex-dependent effects of pathogenic *GRIN* variants, particularly in the study of NDDs, which exhibit male bias in prevalence rates (May et al., 2019; Santos et al., 2022; Bölte et al., 2023; Cruz et al., 2025). The sexual dimorphism of behavioral phenotypes identified in *Grin1*^Q536R/+^ mice provide an opportunity to explore the biological underpinnings of sex differences in individuals with NDDs. *Grin1*^Q536R/+^ mice displayed additional behavioral alterations in social, cognitive, and seizure domains during adulthood, which is in line with the reported clinical characteristics of the patient.

Our study of older adult *Grin1*^Q536R/+^ mice revealed the progression of behavioural phenotypes, where in older animals (unlike adult mice) both male and female variant mice displayed hyperactivity. This is similar to the locomotor phenotypes of other *Grin1* mutant mice, where sexual dimorphism dissipated with age in *Grin1*KD mice (Milenkovic et al., 2014) and *Grin1*^Y647S/+^ mice (Sullivan et al., 2024). These findings reinforce that behavioural alterations imparted by *Grin1* gene variation can evolve throughout adulthood, with certain domains showing potential for improvement with age.

Given the broad distribution of variants across the gene and their diverse impact on NMDAR function, no single model can capture the full spectrum of the pathophysiology, underscoring the need for multiple exemplar models (Benke et al., 2021). Indeed, both patient-derived and experimentally introduced *Grin1* variations have been modeled *in vivo*, which display both overlapping and distinct phenotypes, highlighting the value of variant-targeted approaches (Mohn et al., 1999; Lee, 2023; Sullivan et al., 2024). In this study, the *Grin1*^Q536R/+^ mouse model presents notably milder molecular and behavioral abnormalities compared to existing *Grin1* mouse models, such as the *Grin1*KD, *Grin1*^Y647S/+^, and *Grin1*^G827R/+^ mice (Mohn et al., 1999; Lee, 2023; Sullivan et al., 2024). Importantly, this heterogeneity in *Grin1* mouse models reflects the broad clinical spectrum of *GRIN* disorder. Therefore, including mouse models with mild phenotypes (such as the *Grin1*^Q536R/+^ mice) as part of the exemplar models is critical for a more comprehensive representation of *GRIN* disorder, supporting precision medicine for patients with mild symptoms and investigation of subtle receptor dysfunction and compensatory mechanisms.

In this work, we have contrasted the symptoms associated with a novel genetic variant of the *GRIN1* gene in a patient and in a line of transgenic mice. There are striking parallels in terms of motor outputs and cognition, as well as an overall similarity in the symptom severity. Together with our prior work on *GRIN1* Y647S heterozygous variant, associated with a more severe outcome (Sullivan et al., 2024), we show the potential for many facets of human *GRIN1*-NDDs to be recapitulated in mice. Such mouse-patient dyads give insight into the synapse- and circuit- level disruption of brain function and permit investigation of novel treatment regimes.

## Acknowledgements

We thank Dr. Dawn E. Watkins-Chow and Dr. William J. Pavan from the National Human Genome Research Institute at the National Institutes of Health for their technical assistance in generating the mouse model. The work was supported in part by the Government of Canada’s New Frontiers in Research Fund (NFRF), by Canadian Institutes of Health Research (CIHR) Project Grants awarded to A.J.R. (169153) and E.K.L., CIHR Foundation Grant to G.L.C. (154276), and Simons Foundation Autism Research Initiative (A.J.R. and G.L.C.). Funding and support were also provided by CureGRIN Research Foundation (A.J.R.), the Shireen and Edna Marcus Foundation (A.J.R.) and Dani Reiss Family Foundation (Neurodegeneration and Aging Research Program, G.L.C.). G.L.C. holds the Krembil Family Chair in Alzheimer’s Research.

## Author contributions

M.T.S., Y.Y., P.T., Q.P., T.V.L., S.V., P.S.B.F., J.G., R.E.M., E.K.L., R.P.B., A.S., G.L.C., and A.J.R. designed research; M.T.S., Y.Y., P.T., Q.P., T.V.L., S.V., P.S.B.F., E.F., and W.H. performed research; M.T.S., Y.Y., P.T., Q.P., T.V.L., S.V., and P.S.B.F. analyzed data; M.T.S., Y.Y., and L.V. wrote the first draft of the paper; Y.Y., L.V., P.T., Q.P., P.S.B. F., E.K.L., and A.J.R. edited the paper; Y.Y., L.V., and A.J.R. wrote the paper.

## Supplementary materials

### Materials and Methods

**Genotyping primers**

Primers for WT allele:

5’-AGCCCTTCAAGTACCAG-3’(forward),

5’-CCTCCTCGCTGTTCACCTTAAATC-3’(reverse).

Primers for Q536R variant allele:

5’-AGCCCTTCAAGTACCGC-3’(forward),

5’-CCTCCTCGCTGTTCACCTTAAATC-3’(reverse).

Primers for thrombomodulin (control):

5’-CCAGGCTCTTACTCCTGTAT-3’(forward), 5’-TGGCACTGAAACTCGCAGTT-3’(reverse).

**RT-qPCR primers:**

Primers for mouse *Grin1* (targeting both wildtype and variant transcripts):

5’-ACACCAACATCTGGAAGACAG-3’ (forward),

5’-CAGTCACTCCATCTGCATACTT-3’ (reverse).

Primers for PGK (housekeeping):

5’-GGCCTTTCGACCTCACGGTGT-3’ (forward),

5’-GTCCACCCTCATCACGACCCG-3’ (reverse).

---

**Figure S1.**
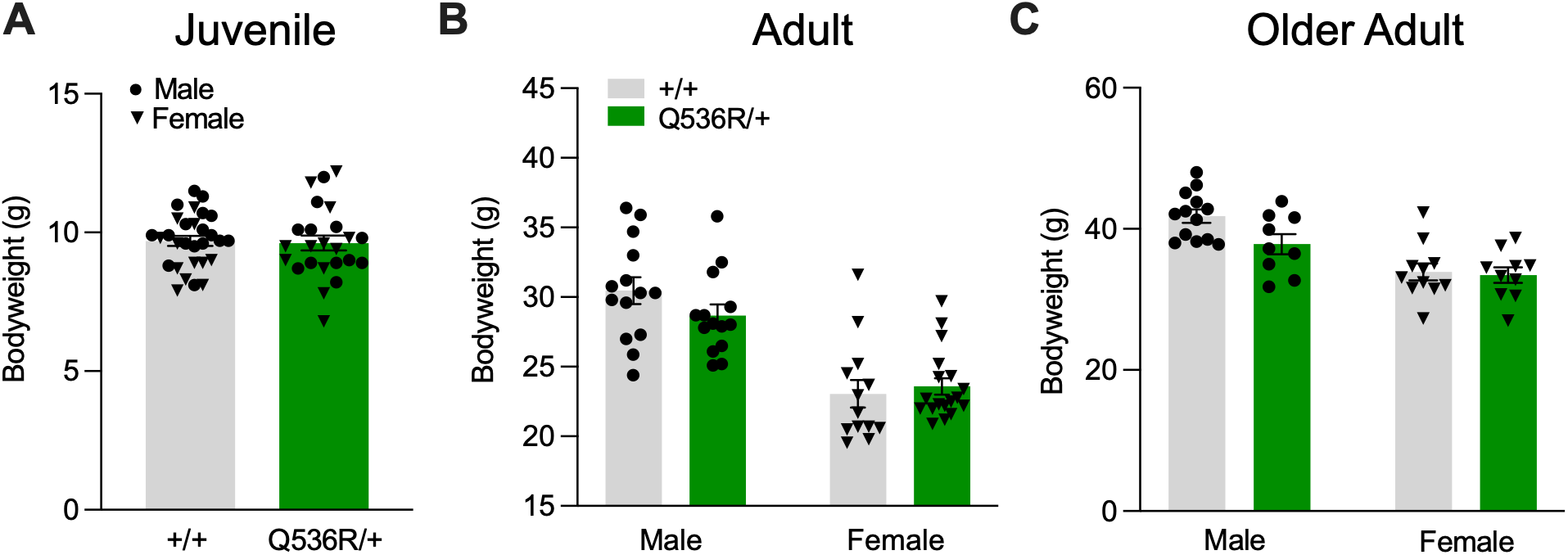
Bodyweight of *Grin1*^Q536R/+^ mice in juvenile, adult and older adult mice. *Grin1*^Q536R/+^ mice do not show changes in bodyweight relative to wildtype mice in **(A)** juvenile (postnatal day 21) (+/+ n = 29, M = 17, F = 12; Q536R/+ n = 24, M = 12, F = 12; p = 0.8182), during **(B)** adulthood (3-5 months) (+/+ n = 27, M = 14, F = 13; Q536R/+ n = 32, M = 14, F = 18; two-way ANOVA: main effect of sex: F[1,55] = 57.44, p < 0.0001) or into **(C)** extended age (9-13 months) (+/+ n = 24, M = 13, F = 11; Q536R/+ n = 19, M = 9, F = 10; two-way ANOVA: main effect of sex: F[1,39] = 28.54, p < 0.0001). Data are expressed as mean ± SEM. Males and females are represented by circle and triangle symbols, respectively.

**Figure S2.**
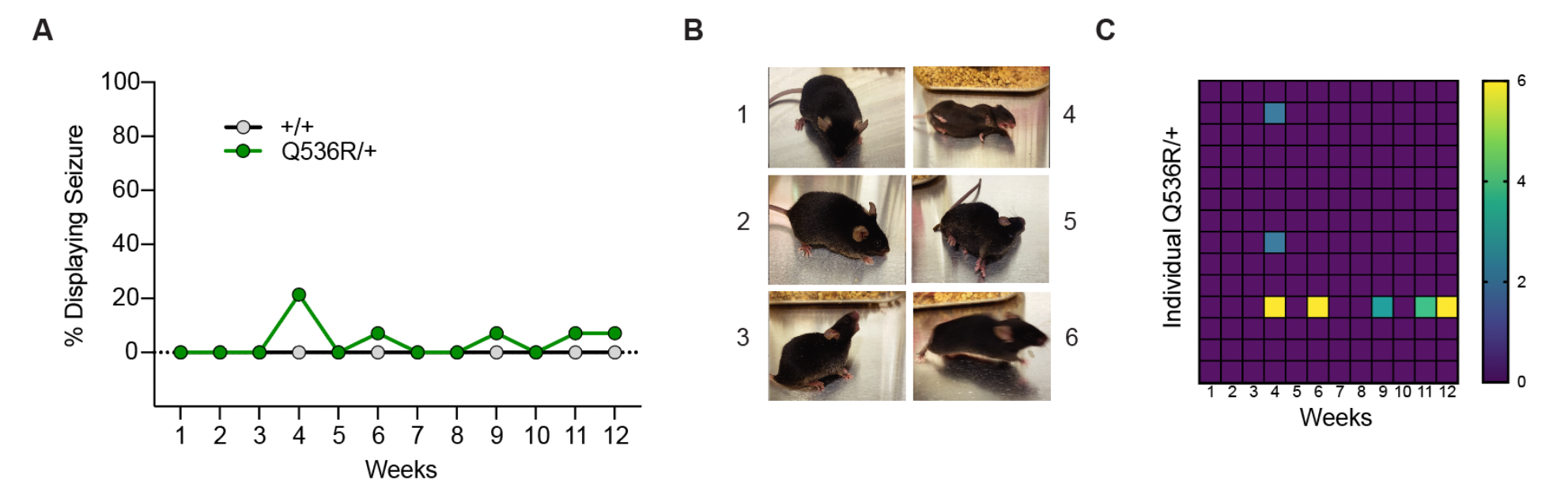
Spontaneous seizure phenotype of adult *Grin1*^Q536R/+^ mice. **(A)** Percentage of +/+ and Q536R/+ mice (age of 22 -29 weeks at Week 1) displaying visually identifiable seizures during weekly handling over 12 weeks (sexes combined: +/+ n = 15, Q536R/+ n = 14). **(B)** Representative photographs of each category of convulsion severity observed during weekly handling as described in **Table S1**. **(C)** Heat map of convulsion severity observed in individual Q536R/+ mice over the course of 12 weeks (n = 14).

**Table S1.** Spontaneous Seizure Severity Scale (Racine scale).

| Seizure Score | Description |
| --- | --- |
| 0 | No change in behaviour |
| 1 | Sudden behavioural arrest, motionless staring |
| 2 | Facial automatisms (whisker trembling, twitching of eyes/ears), minor myoclonic body jerks |
| 3 | Full myoclonic body jerks, head nodding toward back, teeth chattering, drooling, forelimb clonus/clenching while remaining upright, Straub tail |
| 4 | Momentary/partial loss of posture, regaining upright position slowly |
| 5 | Tonic clonic convulsion with loss of righting reflex (mouse completely on its side), forelimb extension |
| 6 | Bouts of jumping/backing up, upright & excessive grooming |

**Table S2.** Current treatment regimen of the patient carrying the Q536R/+ variant.

| Medication | Dosage | Indication | Frequency | Notes |
| --- | --- | --- | --- | --- |
| Atomoxetine | 25 mg | Cognitive Abilities (attention) | Once daily | Started taking recently after switching from Methylphenidate , effectiveness is yet to be determined |
| Valproate | 250 mg | Epilepsy | Twice daily | Seizures have not been observed since re-commencing at age 8. |

